# Tailoring confocal microscopy for in-cell photophysiology studies

**DOI:** 10.1101/2022.11.08.515612

**Authors:** Mattia Storti, Haythem Hsine, Clarisse Uwizeye, Olivier Bastien, Daniel Yee, Fabien Chevalier, Cécile Giustini, Daniel Béal, Johan Decelle, Gilles Curien, Dimitri Tolleter, Giovanni Finazzi

## Abstract

Photoautotrophs environmental responses have been extensively studied at the organism and ecosystem level. However, less is known about their photosynthesis at the single cell level. This information is needed to understand photosynthetic acclimation processes, as light changes as it penetrates cells, layers of cells or organs. Furthermore, cells within the same tissue may behave differently, being at different developmental/physiological stages. Here we describe a new approach for single-cell and subcellular photophysiology based on the customisation of confocal microscopy to assess chlorophyll fluorescence quenching by the saturation pulse method. We exploit this setup to: i. reassess the specialisation of photosynthetic activities in developing tissues of non-vascular plants; ii. identify a specific subpopulation of phytoplankton cells in marine photosymbiosis, which are consolidating metabolic connections with their animal hosts, and iii. testify to the link between light penetration and photoprotection responses inside the different tissues that constitute a plant leaf anatomy.

**Motivation:** Visualising photosynthetic responses in 3D is essential for understanding most acclimation processes, as light changes within photosynthetic tissues as it penetrates the absorbing/diffusing layers of the cells. To achieve this goal, we developed a new imaging workflow merging confocal microscopy and saturating pulse chlorophyll fluorescence detection. This method applies to samples characterised by increasing complexity and its simplicity will contribute to its widespread use in plant and microalgae photoacclimation studies.

## Introduction

Photosynthesis is a major bioenergetic process in the biosphere. It feeds most of the food chains on Earth and is responsible for substantial sequestration of CO_2_ via the biological pump. The efficiency and regulation of this process is usually assessed *in vivo* by measuring chlorophyll (Chl) fluorescence (Baker, 2008), i.e. the fraction of absorbed light that is re-emitted in the near-infrared region of the spectrum. Chl fluorescence can be analysed at different scales: from the cellular level using microscopes to organs/organisms using IR cameras or even at larger scales (ecosystems/planetary) using satellites (Meroni et al., 2009; Ni et al., 2019; Yang et al., 2015). Those approaches all suffer from a similar limitation, namely the detection of images in two dimensions only. However, significant changes are expected within the 3D volume of phototrophs, because the colour and intensity of light vary according to its penetration into the absorbing/diffusing layers of photosynthetic cells (Ptushenko et al., 2020). So far, theoretical approaches were used to extrapolate data obtained from a surface (leaf, canopy or ocean) to a volume, by modelling light penetration (Evans, 1999). In a few cases, the responses of a photosynthetic tissue (e.g. a leaf) to different colours of light (blue, green and red) have been compared to evince the effect of different light penetrations on photosynthesis (Qi et al., 2003; Rappaport et al., 2007; Terashima et al., 2009). Both approaches have limitations as they do not rely on direct assessment of photosynthesis inside an intact photosynthetic tissue/organ. Moreover, monitoring fluorescence changes in 2D plan does not allow to detect fluorescence of single plastid, because these organelles move inside cells (Cazzaniga et al., 2013; Sato et al., 2001; Wada et al., 2003) do not allow to monitor single plastid fluorescence kinetics usually acquired on a plane. To overcome this difficulty, we have explored the potential of a new Chl fluorescence imaging approach that combines the spatial resolution of a confocal microscope with the reliability of the saturation pulse method (Schreiber, 2004). The latter approach has been particularly successful in assessing relevant photosynthetic parameters (the quantum yield of Photosystem (PS) II in the dark -Fv/Fm- and in the light -ΦPSII-; the thermal dissipation of excess excitation energy -NPQ- (Maxwell & Johnson, 2000) to study CO_2_ assimilation capacity, plant acclimation to the environment and stress responses (Ogawa et al., 2017). We show that the 3D saturating pulse confocal setup provides unique physiological information concerning photoprotection in biological samples characterised by increasing complexity: *i*. heterogeneous responses of single chloroplast/cell in mosses, which contain multicellular and partially differentiated tissues. *ii*. complex relationships in photosymbiosis between eukaryotic host-cells and symbiotic microalgae at different developmental stages. *iii*. the link between leaf architecture and photoprotection in vascular plants.

## Results

### Combining a saturating pulse method with a confocal microscope

To develop a 3D saturating pulse confocal (hereafter 3D-Pulse fluorimeter), we equipped a Zeiss LSM900 inverted confocal microscope with an additional red light LED source (λ = 630 nm, Full Width-Half Maximum 18 nm) placed in front of the microscope objective (Fig. 1A). The LEDs deliver short, intense pulses (2000 μmol photons m^-2^ s^-1^) to saturate PSII and thus achieve maximum Chl fluorescence emission, Fm (Butler, 1978; Maxwell & Johnson, 2000). The LEDs also provide continuous actinic light of adjustable intensity (actinic light), to achieve steady-state (Fs) fluorescence (Fig. 1B). Both the pulses and the continuous light delivered by the LED array are operated by a home-built control box. The blue laser (λ = 488 nm) of the confocal is used as the ‘measuring light’ in the saturation pulse method (Schreiber, 2004) to image Chl fluorescence (Fig. 1A), leading to a parameter hereafter called F’. This parameter is close to the F0 parameter used in the saturating pulse method. (Schreiber, 2004). Both the blue laser, the acquisition (on a predefined Z-stack) and the LED control box are controlled by the confocal software Zen (version 3.0), through the “experiment designer” routine. We used the Fiji software (Schindelin et al., 2012) to analyse the data sets and treated the images as follows. 3D time series acquisitions (xyzt) were converted to 2D images (xyt) in a way that preserves original fluorescence values (‘sum slices’ function). The latter can be calculated for every time point measuring the ‘mean gray value’ of Regions Of Interest (ROIs, chloroplasts, cells and tissues) and subtracting the background fluorescence (i.e. the signal measured in an empty ROI located near the measurement region). Indeed, background fluorescence is also affected by external light source (Figure S1). When needed, image segmentation was done with the 3D Slicer software (Uwizeye, et al., 2021), to generate 3D models that gave information about the plastids volume. We calculated photosynthetic parameters from fluorescence values with the Origin software (Microcal, USA).

**Figure 1.**
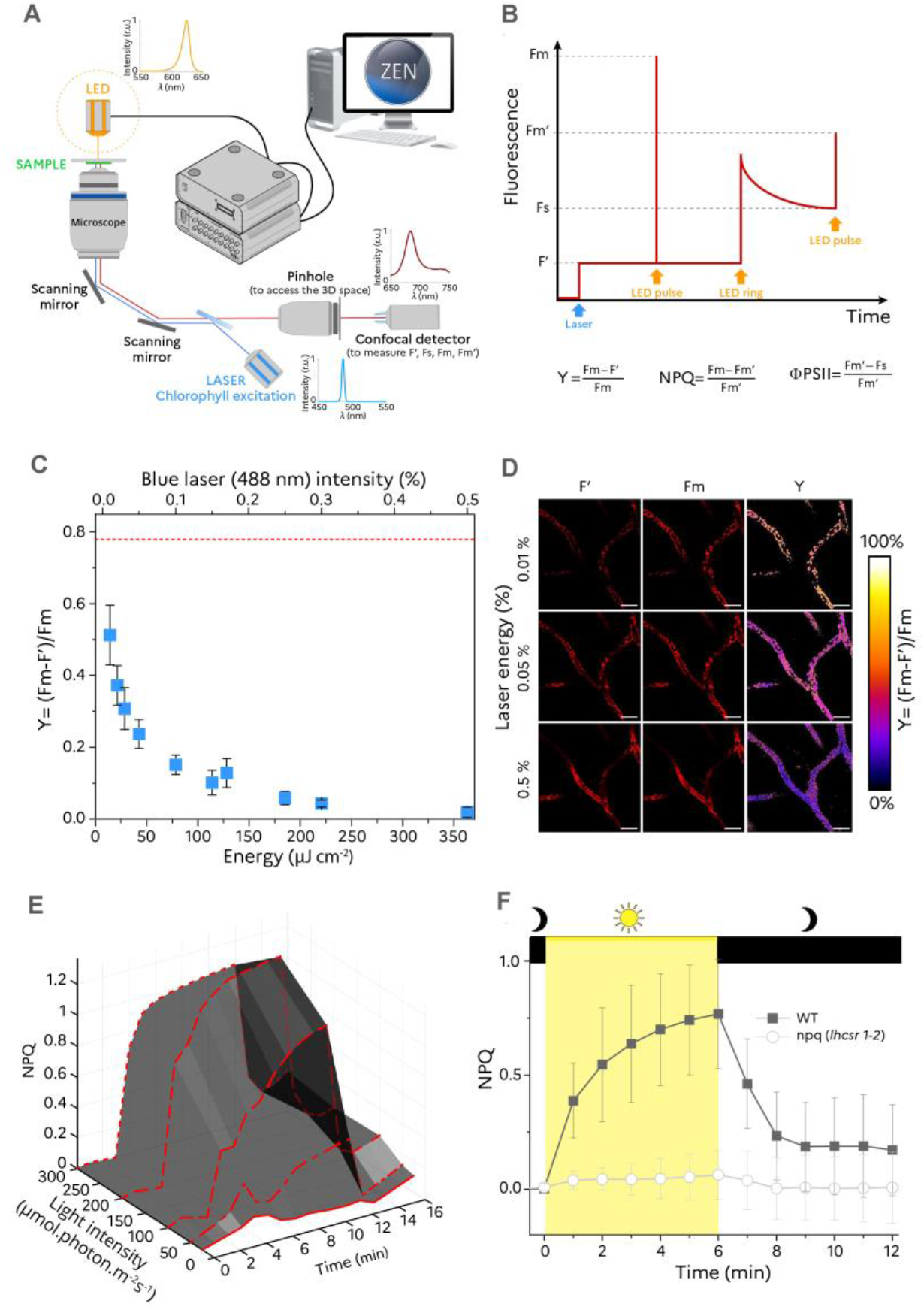
3D-Pulse fluorimeter imaging setup. (A), The customised confocal microscope includes an orange LED array controlled by the Zen software via the SVB1 module plus a homemade controller to deliver actinic light and saturating pulses. (B), Switching on and off the light pulses via the ‘experimental design’ routine (and manual switching on and off the actinic light) allows evaluating relevant photosynthetic parameters. (C), The apparent quantum yield of photosystem II (Fm-F’)/Fm is lower that values measured with a standard Chl fluorescence imaging camera (red dashed line), and decreases as a function of the energy of the confocal laser power (blue squares, N=7 cells average ± s.d.). (D), Representative Chl fluorescence images used to calculate Y, scale bar: 50 μm. (E), NPQ changes as a function of the light intensity (25, 50, 100, 200 and 500 μmols photons m^-2^ s^-1^). Representative traces of an experiment repeated 5 times with similar results. (F), NPQ features in a WT (solid symbols, average of 108 cells ± s.d.) and mutant (open symbols, average of 63 cells ± s.d.) with downregulated NPQ capacity. Dark box: actinic light off; yellow box: actinic light (500 μmols photons m^-2^ s^-1^) on.

We first validated the 3D-Pulse fluorimeter on a photosynthetic organism having a relatively simple structure, the juvenile gametophyte (protonema) of the moss *Physcomitrium patens*. We noticed that the maximum photosynthetic capacity, here indicated by the PSII related parameter Y = (Fm-F’)/Fm, decreased when the laser intensity increased. The Y decrease reflects the actinic effect of blue laser itself, which increases the F’ parameter. At 0.3% laser intensity (i. e. 220 μJ.cm^-2^, Fig. 1C), the F’, obtained in the presence of the blue laser illumination alone, reached the same level as Fm, which is the fluorescence intensity obtained by concomitant illumination with the blue laser and the saturating red pulse. This finding suggests that the blue laser alone, which is localised over a very small area, saturates photosynthesis (inducing Fm) even at relatively low intensities. We exploited this possibility to measure Fm and Fm’ without the saturating pulse protocol, and therefore to calculate NPQ, which is readily estimated from these two parameters (Fig. 1B). On the other hand, the relative PSII yield can be evaluated using a subsaturating laser intensities (Fig. 1D).

The 3D-Pulse fluorimeter was fast enough to measure the kinetics of NPQ onset (when the actinic light was switched on) and its relaxation in the dark (Fig. 1E, Supplementary Videos 1-4). We could also detect changes in NPQ amplitude as a function of actinic light intensity, as well as a transient NPQ during exposure of the dark-adapted protonema filaments to low light intensity (Fig 1E, dash and dotted line). This transient NPQ reflects the link between activation of CO_2_ assimilation and photoprotective responses: at the beginning of illumination, when CO_2_ assimilation is largely inactive, part of the absorbed light is dissipated. Conversely, most of the absorbed photons are drained to CO_2_ assimilation in steady state (when the Calvin Benson Bassham cycle is fully active) and thus NPQ disappears (Finazzi et al., 2004). Finally, we could easily differentiate NPQ features of a WT and a mutant strain with reduced NPQ capacity (due to knocking out of the NPQ effector proteins LHCSR1 and LHCSR2 ((Gerotto et al., 2012), Fig. 1F). We noticed that NPQ was lower in the 3D-Pulse fluorimeter than in a conventional 2D imaging fluorimeter equipped with the same light sources and intensities (Figure S2). This difference could be caused by the low amount of actinic light delivered inside the tissue (where the 3D-Pulse fluorimeter measures), when compared to its surface, where conventional 2D imaging quantifies NPQ.

### Photoprotective responses in non-vascular plants

We explored the possibilities offered by 3D-Pulse fluorimeter to study cellular and subcellular NPQ responses in the two types of cell that constitute the protonema of *P. patens* (Cove, 2005): the caulonema and the chloronema. The former has longitudinally elongated cells involved in propagation and nutrient acquisition, the latter has chloroplast-rich cells, usually considered as the photosynthetic part of the moss protonema (Fig. 2A) (Cove, 2005). As plastids move inside the cell (Sato et al., 2001; Yamashita et al., 2011), they tend to leave the field of observation in a conventional microscope during the relatively long time required for NPQ development (Supplementary Video 1). Instead, we could follow plastid responses within the entire volume of caulonema and chloronema cells with the 3D-Pulse setup, (Supplementary Videos 2-5), visualise Chl fluorescence (Fig. 2B), cell fraction occupancy (Fig. 2C) and quantify NPQ capacity (Fig. 2D) of single cells and plastids. A principal component analysis (PCA) of 175 mutant and WT cells allowed to distinguish four classes (Fig. 2E, Supplementary Tables 1-3): the WT (circles) and the NPQ mutant *lhcsr1/2* (triangles) were separated based on their NPQ capacity, while the two cell types (black: chloronema; grey: caulonema) could be differentiated because of their plastid cell density. A more refined analysis (WT, Fig. 2F, see also Figure S3 for the *lhcsr1/2* mutant) revealed subtle heterogeneity in NPQ responses in both caulonema and chloronema, which we could interpret based on 3D imaging. We identified heterogeneous NPQ responses at the cell level (Fig. 2G), which account for most of the above-mentioned heterogeneity. Conversely, single plastids (Figure S4) within a given cell behave homogeneously (Fig. 2H): cells with high photoprotective responses contain plastids with high NPQ capacity, while cells with low photoprotection have plastids with low NPQ. Such variability in NPQ responses at the cell level probably reflects the different physiological state of cells that continuously regenerate during protonema development. While it is relatively easy to distinguish caulonema from chloronema in the complex matrix of the protonema, it is difficult to attribute more specific cellular characteristics (such as their age), which certainly have an impact on photosynthetic behaviour.

**Figure 2:**
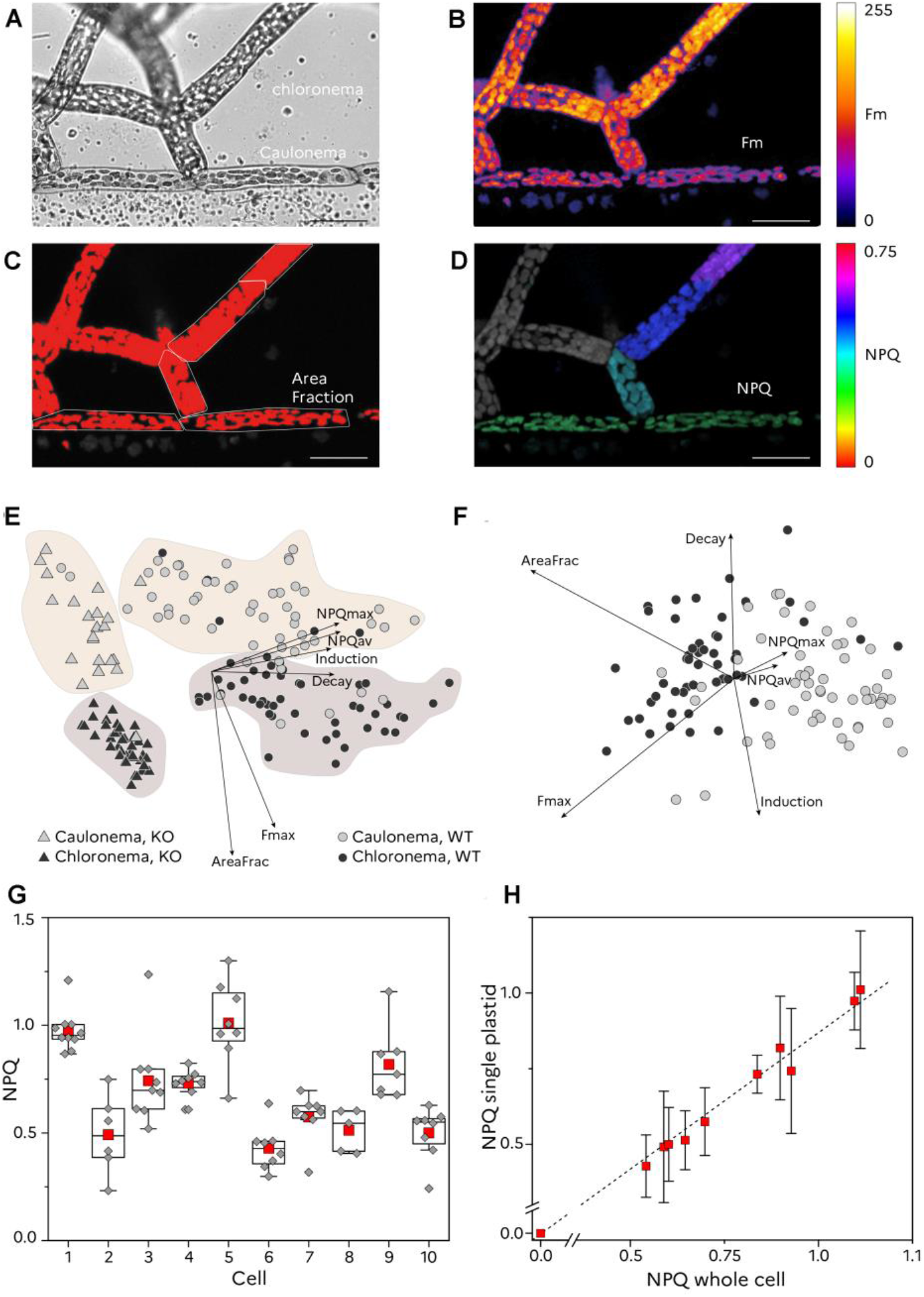
NPQ features in growing tissues of the moss *Physcomitrium patens*. (A), bright field image. (B), false colour image of chlorophyll fluorescence. (C), fraction of the cells occupied by plastids (red). (D), false colour image of NPQ in caulonema and chloronema cells of *P. patens* protonema, scale bar 50 μm. (E,F), Principal Component Analysis of 175 cells (black circles: chloronema wt, black triangles: chloronema mutant, grey circles: caulonema wt, grey triangles: caulonema mutant). (E): First and second components; (F): second and third components for WT cells. The first two components represent roughly 88% of the variance, while the first three components represent more than 94% of the variance (Supplementary Table 1). (G), NPQ is heterogeneous in *P. patens* cells. Red squares: mean of NPQ values of all plastids inside the same cell (grey diamonds; boxes: P25 and P75; whiskers: outliers, black line: median. (H), Plastid vs cells NPQ relationship reveals that plastids (average of 5-10 ± s.d.) behave homogeneously inside a given cell.

### Probing heterogeneous photosynthetic activity inside a complex, photosymbiotic organism

We further investigated the ability of the 3D-Pulse fluorimeter to link NPQ responses to different physiological states focusing on photosymbiosis, a common lifestyle in oceanic plankton between symbiotic microalgae and unicellular eukaryotic hosts (Biard et al., 2016; de Vargas et al., 2015; Decelle et al., 2012; Guidi et al., 2016; Michaels et al., 1995). Inside hosts (radiolarians), symbiotic microalgae (the haptophyte *Phaeocystis cordata*, Fig. 3A) undergo progressive morphological and metabolic changes. Therefore, a single host cell contains a mix of newly engulfed/small symbionts with two plastids, and of larger (presumably older) ones, with up to 60 plastids as revealed by focused ion beam scanning electron microscopy imaging (FIB-SEM) (Decelle et al., 2019; Uwizeye, et al., 2021). Using confocal microscopy, we confirmed the algal heterogeneity in terms of cell volume occupied by plastids (Uwizeye, et al., 2021) (Fig. 3B, histogram). Moreover, we were also able to extract quantitative photosynthetic characteristics of individual microalgae inside the host upon segmentation and 3D reconstruction of their Chl fluorescence emission (Fig. 3B).

**Figure 3:**
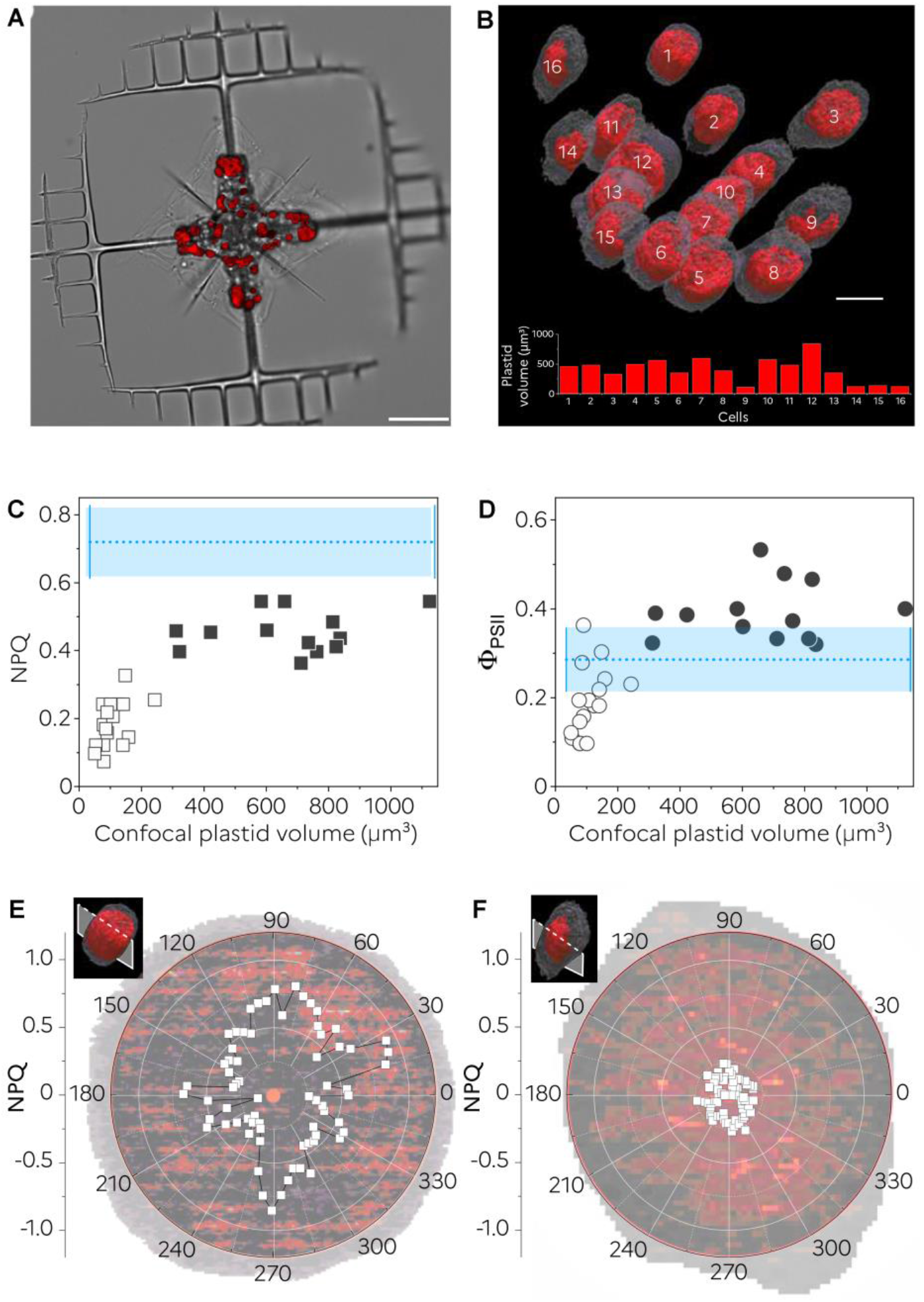
photosynthetic features in a planktonic photosymbiosis. (A) Bright field (grey) and Chl fluorescence (red) images overlaid from a host acantharian cell harbouring symbiotic microalgae (the haptophyte *Phaeocystis*), Bar: 50 μm. (B) 3D reconstruction of plastid chlorophyll fluorescence inside 16 microalgae (top) reveals large differences in their plastid volumes (bottom), Bar: 10 μm. (C,D) single cell analysis reveals the existence of two symbiont populations having different photoprotective responses (NPQ) and photosynthesis (ΦPSII). Blue dotted lines: NPQ and ΦPSII in free living *P. cordata* cells (N = 15 ± s.d.-blue box) (E,F) radar plot of the NPQ in large (E) and small (F) symbiotic microalgae. NPQ was calculated from images acquired in the dark and after exposure to actinic light (500 μmols photons m^-2^ s^-1^) for 15 minutes. Representative traces of an experiment performed on 8 cells (e) and 5 cells (f) respectively (Figure S5).

Relating plastid volume heterogeneity to single-cell NPQ responses revealed the existence of two symbiont populations: small algae (open symbols) exhibited lower NPQ (Fig. 3C) and photosynthetic activity (assessed by the ΦPSII parameter, Fig. 3D) than free-living cells measured with the same 3D-Pulse fluorimeter (Fig. 3C,D blue dotted line). Conversely, larger symbionts (solid symbols) had a different trend: their NPQ was always lower, while ΦPSII was higher than in free-living cells. We interpret the lower NPQ but higher ΦPSII of larger symbionts as a signature of enhanced photosynthetic performance upon transformation of the alga inside the host, as previously reported (Decelle et al., 2019).

The first symbiont population (small algae) has not been reported so far in photosymbiosis, likely because it represents a relatively small fraction of the symbionthic cells, difficult to observe with conventional chlorophyll fluorescence imaging setups. Their photosynthetic features (concomitant decrease of photosynthesis and NPQ) remind the ones observed in the diatom *Phaodactylum tricornutum* (Allen et al., 2008) and the green alga *Chlamydomonas reinhardtii* (Naumann et al., 2007) upon mineral nutrient (Fe) limitation. We propose that this particular population comprises microalgae in the process of adapting to the trophic environment provided by the host (see Discussion).

To take the analysis a step further, we investigated NPQ features at the subcellular level. However, it was difficult to track Phaeocystis plastids separately based on confocal images, because they are around 10 times smaller than in plants and *P. patens* (Crumpton-Taylor et al., 2012; Takemura et al., 2017), and very densely packed (Uwizeye, et al., 2021). Therefore, we developed an alternative approach: we started from 2D confocal slices, where we approximated the fluorescence emitting area with a circle. Inside this area, we scanned the Chl fluorescence intensity values along a single radius in the dark and after 10 minutes of illumination, to calculate Fm, Fm’ and NPQ ((Fm-Fm’)/Fm’). We repeated this calculation every 5 degrees, to infer the homogeneous/heterogeneous NPQ responses of inside the cells. We found very heterogeneous NPQ values in both large (Fig. 3E) and small (Fig. 3F) symbionts (see also Figure S5). This result suggests that, in contrast to mosses, plastids of photosymbiotic cells develop NPQ responses in a rather heterogeneous manner. In contrast, homogeneous plastid responses, such as those reported above in P. patens, should result in relatively constant NPQ values for each angle measured

### Vascular plants NPQ is regulated by light channelling throughout anatomically diverse leaf architectures

Finally, we exploited the 3D-Pulse fluorimeter to investigate fluorescence responses in highly complex photosynthetic architectures: vascular plant leaves. These highly efficient machineries are composed of millions of cells, receiving variable light intensity depending on their position inside the organ. Cells on the surface receive more photons than cells inside the leaf, resulting in a light gradient (Wuyts et al., 2012). The light gradient in turn leads to a different extent of saturation of photosynthesis, which we inferred via the amount of light in excess dissipated via NPQ. Thanks to high resolution of the 3D-Pulse fluorimeter, and the penetration of the blue laser, we could measure NPQ in mesophyll cells around 80 μm under the epidermis (vertical bars with blue arrows in Fig. 4) of a leaf exposed to the red actinic light on the opposite side of the detection. In a monocotyledon (*Plantago lanceolata*), we recorded similar NPQ responses on the adaxial and abaxial sides (Fig. 4A,C) consistent with the observation of a symmetrical leaf morphology (Fig. 4B), and thus of similar light gradients in both directions (Figure S6).

**Figure 4:**
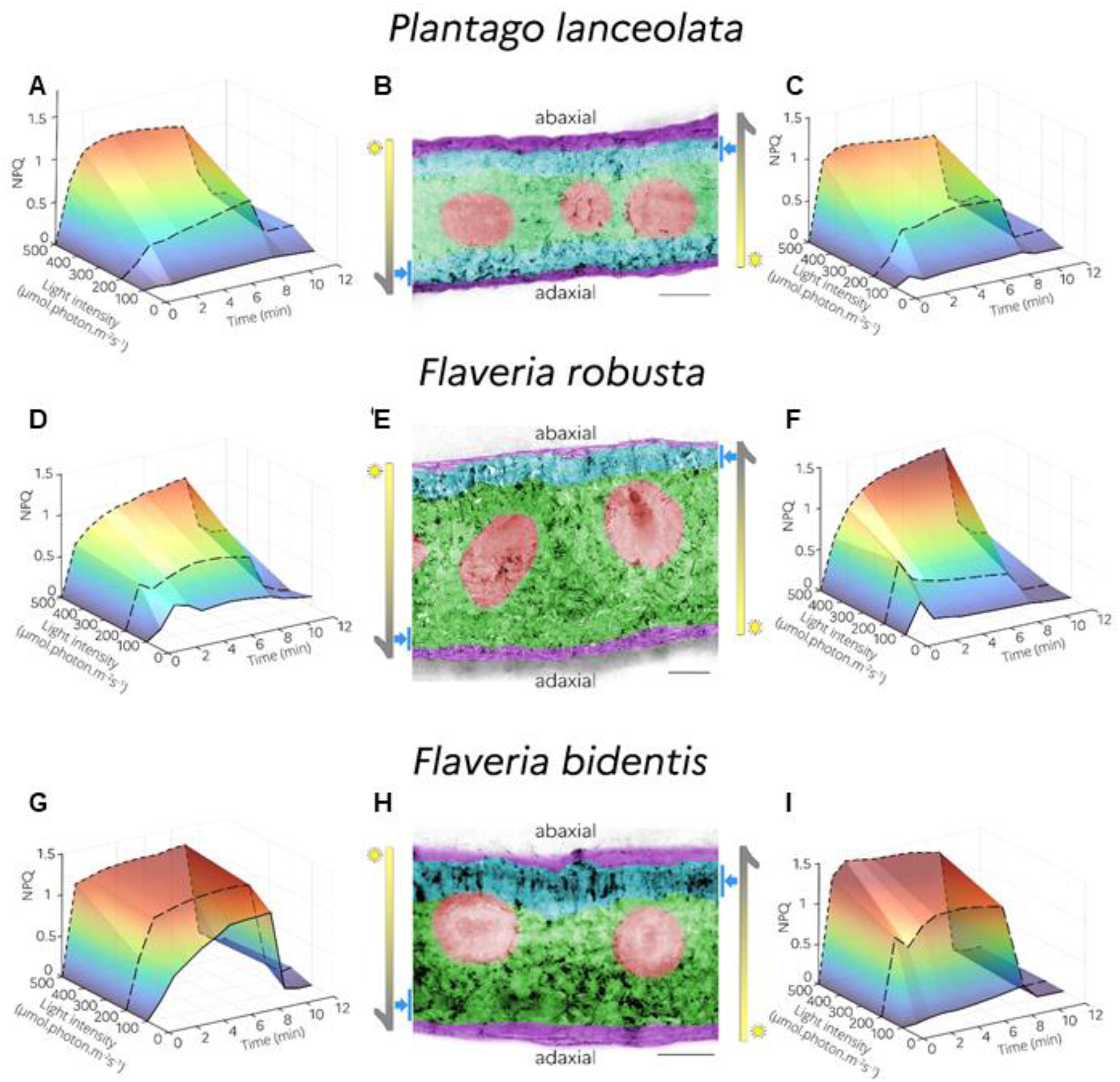
NPQ features are modulated by leaves architectures. NPQ measured upon illumination in the adaxial to abaxial side (top to bottom –panels (A, D and G) and abaxial to adaxial (bottom to top – panels (C, F and I) for three leaves with different anatomies. Representative picture of an experiment repeated 3-7 times with similar results. (B), (E), (H), False colours representation of the different leaf tissues: purple: epidermis; blue: palisadic parenchyma; green: spongy parenchyma; red: vascular tissue. Vertical bars with blue arrows represent the region imaged in NPQ experiments. Bar: 100 μm

Conversely, we observed heterogeneous NPQ responses in dicotyledons (*Flaveria robusta* and *Flaveria bidentis*), which harbour an asymmetric mesophyll organisation (Fig. 4E,H): on the adaxial face, the palisading parenchyma (blue) under the epidermis (purple), is made up of elongated photosynthetic cells arranged perpendicular to the leaf surface. On the opposite abaxial side, this layer is replaced by the lacunar parenchyma (green), which occupies a large part of the leaf surrounding the vascular tissues (red) and consists of more irregular cells and large intercellular spaces to promote gas circulation and storage.

In both Flaveria species, NPQ at non-saturating light intensities was higher when actinic light was provided from the adaxial side to abaxial side (top to bottom, Fig. 4D,G) compared to actinic light provided from abaxial to adaxial side (bottom to top, Fig. 4F,I). By combining our 3D-Pulse fluorimeter derived findings with a more ‘classic’ approach (Evans, 1999) (measuring light penetration gradients inside a leaf, Figure S6) we interpret these data as follows: thanks to channelling through the parallel cell layers of the palisade mesophylls, light better penetrates the leaf in the adaxial to abaxial direction (Figure S6). Therefore, it saturates photosynthesis at a lower photon density and induce high photoprotective responses (NPQ) (Figure S7). Conversely, illumination in the abaxial to adaxial direction appears to favour light scattering from randomly oriented spongy cells. This phenomenon decreases photon penetration, lowers excess light, and therefore NPQ. Consistent with this conclusion, we observed that the dissymmetry of the NPQ was exacerbated in *F. bidentis*, where the steepness of the light gradient is higher due to a reduced thickness of the leaf (Figure S6). In agreement with this notion, differences in NPQ were erased when the light was increased to 500 μmols photons m^-2^ s^-1^, an intensity that should oversaturate photosynthesis irrespectively of the direction of illumination. Hence, our method allowed to characterise *in situ*, without the use of models or assumptions, the effect of differential light penetration on photosynthesis.

## Discussion

In this work, we show that imaging Chl a fluorescence with a 3D saturating pulse confocal setup is well suited to image photosynthetic responses in 3 dimensions. Previous attempts to use a confocal microscope to measure photosynthesis (Omasa et al., 2009; Tseng & Chu, 2017) suffer from the limitations of using the confocal laser as the ‘measuring light’, the ‘actinic light’ and the ‘saturating pulse’ of the Saturating pulse approach. This choice implies that two different lights cannot be provided as the same time (unlike in the saturating pulse approach), thus hampering the accuracy in the determination of the photosynthetic parameters which and the number of possible applications (e.g. discriminating the stomata from epidermis in plant leaves, assessing single plastid fluorescence transients (Omasa et al., 2009; Tseng & Chu, 2017)). Conversely, our setup combines the sensitivity and flexibility of the Saturation Pulse Method with the 3D spatial resolution of confocal microscopy to investigate photosynthesis *in vivo* at the organ, cell and subcellular levels.

Thanks to this approach, we obtained original information on photosynthesis in organisms characterised by increasing biological complexity. Our analysis of non-vascular plants revises the classical concept that chloronema cells alone contribute to carbon fixation within the protonema, while caulonema cells are mostly involved in nutrient acquisition and tissue propagation (Thelander, 2005; Xiao et al., 2011). Indeed, while the two cell types can easily be distinguished on the basis of the cell fraction occupied by plastids (Fig 2F, AreaFrac parameter) and total Chl fluorescence (Fig 2F, Fmax parameter), there is almost no difference when photosynthetic parameters (Fig 2F, NPQmax, NPQav, decay and induction parameters) are considered. Thus, we conclude that the plastids of both cell types have similar photosynthetic performance, and that the higher photosynthetic activity of chloronema cells probably stems from their higher plastid content, which makes their photosynthesis easier to follow with conventional chlorophyll fluorescence imaging.

In radiolarians, we identified a particular algal subpopulation characterised by very low photosynthetic performance and photoprotection capacity. In general, these two parameters show complementary responses: low photosynthesis results in high dissipation of excess light in the form of NPQ, whereas high photosynthesis leads to low dissipation of excess light. The latter behaviour is indeed observed in the large cells (Fig 3 C,D), which probably represent ancient algae that have established fully metabolic connections with the host (Uwizeye, et al., 2021). Conversely, the population with low photosynthetic performance is made up of small cells, probably still adapting to the host trophic environment(Uwizeye et al., 2021). Indeed, a concomitant decrease in photosynthesis and photoprotection responses (NPQ) has only been reported in microalgae exposed to mineral nutrient (Fe) limitation (Allen et al., 2008; Naumann et al., 2007). It is tempting to draw a parallel between this population, which likely represents a transient stage in the establishment of photosymbiosis, and the hypothetical early stage of endosymbiosis (Bhattacharya et al., 2007; Cenci et al., 2017; Karkar et al., 2015), in which metabolic connections between cyanobacteria/algae and the animal host must be established, prior to the transfer of genetic information from the phototroph to the animal recipient organism.

Our analysis of leaf photosynthesis supports previous findings that light is differentially channelled through different leaf tissues (Evans, 1999). However, we go one step further, showing that moderate light intensity, which is on average received by most leaves in a tree due to mutual shading, can saturate photosynthesis over the entire leaf section in asymmetric leaves (Fig. 4E and F). This is, however, only true when photons are captured on the abaxial side, where the palisading parenchyma channels them toward the opposite side of the leaf (Fig. 4D and G), the spongy parenchyma. Overall, these results confirm the notion that leaf photosynthesis is largely governed by its anatomical features (e.g. Wright et al., 2004) and further extends it to the cellular level.

### Limitations of Study

Our 3D imaging approach has clear advantages over conventional PAM fluorometers and single-cell pulse-probe microscopes, since it allows: i. disentangling heterogeneous cellular responses within a tissue (Fig. 2); ii. tracking photosynthetic changes during cell development (Fig. 3); iii. assessing the relationship between leaf anatomy (asymmetric or symmetric) and photoprotection (Fig. 4). Its 3D resolution allows to follow the movements of plastids during prolonged exposure to light (Supplementary Videos 1-4) in order to monitor their physiological responses.

However, there is a main difficulty associated with this approach. The choice of the laser light intensity is essential to ensure correct measurements of the photosynthetic parameters. In our case, the maximum photosynthetic capacity (the ‘Y’ parameter in Fig. 1B) decreases as a function of laser intensity (Fig. 1C), as PSII becomes inactive (light saturated). The PSII is completely saturated (i.e. the ‘Y’ parameter goes to 0) at 0.3% of the maximum power, indicating that the laser is too intense. In this case, this difficulty could be alleviated by placing neutral filters between the laser and the sample to decrease the laser power. Other organisms (or setups) may have different responses to light, so an experiment similar to that shown in Fig. 1C is recommended before undertaking NPQ measurements.

In principle similar information could be obtained using a fluorescence Lifetime Imaging Microcope (FLIM) (Iermak et al., 2016; Iwai et al., 2010). This approach relies in a direct measurement of the chlorophyll fluorescence decay lifetimes, and therefore also provides information about photochemical use and thermal dissipation excitation energy. While this approach has the advantage of being insensitive to the relative power of the confocal laser, it requires a dedicated (and expensive) setup, making its use more restrained. Conversely, our extremely affordable setup can in principle be easily connected to any existing confocal microscope, it represents an alternative tool to study photosynthetic acclimation responses at the cellular/subcellular level, plant cell-animal cell interactions, trophic lifestyles, and to monitor plant and microalgal development.

## Methods

### Photosynthetic material

#### Physcomitrium patens

Gransden wild-type (WT) strain and *lhcsr1-lhcsr2* KO (*lhcsr1/2* (Gerotto et al., 2012)) were grown on PpNO3 (3mM Ca(NO_3_)_2_, 1 mM MgSO_4_, 50μM FeSO_4_, 200 μM KH_2_PO_4_ pH 7, traces elements (Ashton et al., 1979)) solidified media (0.8% Plant Agar) overlaid with a cellophane filter. Moss tissue was propagated vegetatively by homogenization through a tissue blender and cultivated in axenic condition at 25°C, 40 μmol photons m^2^ s^-1^ continuous illumination. 10-days-old moss protonema was employed for confocal microscopy measurement.

#### Flaveria robusta

*Flaveria bidentis* and *Plantago lanceolata* plants were grown 3-4 weeks in controlled growth chambers in long day conditions (16h light/8h dark) at a PPF of 100 μmol photons m^−2^ s^−1^. Air temperature was 22°C during the day and 21.0°C at night. Relative humidity was constant at 70% during the day and night. Young fully developed leaves were chosen for experiments.

Symbiotic acantharians harboring intracellular microalgal cells (*Phaeocystis cordata*) were gently collected by towing a plankton net of 150 μm in mesh size with a large cod-end (1 L) for 1-2 min in surface waters (Mediterranean Sea, Villefranche-sur-Mer, France). After collection, individual cells were isolated under a binocular with a micropipette (Decelle et al., 2012). Cells were rapidly transferred to natural seawater and maintained at 20°C and 100 μmol photons m^2^ s^-1^ controlled illumination. Samples were imaged within 24h from sampling time. Cultures of the haptophyte *P. cordata* (the symbiont of Acantharia in the Mediterranean Sea algal (Decelle et al., 2019), strain RCC1383 from the Roscoff Culture Collection) were maintained at 20°C in K5 culture medium at 100 μmol photons m^-2^s^-1.^20°C.

### Sample preparation

*P. patens*. A small portion (∼5mm diameter) of *P. patens* protonema was endorsed on a 20×20 mm coverslip and soaked in 100 μl of water in order to spread the filaments. The coverslip was fixed to the microscopy slide by using a double-side tape of 0.15 mm thickness and finally sealed using VALAP (1:1:1 Vaseline, Lanolin, Paraffin) to prevent water evaporation.

*P. cordata* and *Acantharians* were settled on a microscopy glass bottom dish (Ibidi, Germany) directly on the microscopy plate to avoid media perturbation.

*F. robusta, F. bidentis* and *P. lanceolata*. 5×3 mm sections (longer axis parallel to major leaf veins) were excised using a scalpel. Sections were enclosed between two 20×20mm coverslips allowing to observe both sides of the leaves. A double layer of tape was used to create an enclosure to lodge the leaf section and prevent its crushing. Water was provided to the sample to prevent dehydration during NPQ measurements.

All samples were dark adapted for at least 20 min before confocal microscopy measurements.

### Confocal microscope setup for fluorescence measurement

The Zeiss LSM 900 microscope was equipped with continuous light and pulses of strong actinic light, provided by a LED module located in front of the biological sample. The module contains 4 red LEDs (OSRAM, LA W5AM, λ = 630 nm, Full Width-Half Maximum 18 nm), equipped with a lens to reduce their divergence. The 4 LEDs were oriented at 42°, to focus their light onto the middle of the sample holding slits. The LEDs deliver actinic light, the intensity of which (200 and 500 and 1300 μmols photons m^-2^ s^-1^) is determined by a 3-position switch located on the front face of the control box. Whenever needed, lower actinic radiation levels were obtained by placing a neutral filter (Kodak ND0.3) between the LED array and the sample. This output is connected to the SVB1 Zeiss module, which controls fluorescence acquisition via confocal microscope through the ‘experimental design’ routine of the ZEN software (version 3.0 Blue edition). The same routine also triggers the switching on of saturating pulses (2000 μmols photons m^-2^ s^-1^, duration 1.1 s) to measure of Fm and Fm’. Alternatively, saturating pulses can be switched on manually through a button located on the control box.

Images were acquired with a 20x/0.8 M27 objective (Zeiss) in confocal mode setting with Airyscan detector. A maximum of 40 z-stacks (512 * 512 pixels, 2μm in z between two frames) were acquired at the fastest speed (pixel dwell time 1.03 μs, with an illumination of 633 ms per frame), to minimise light exposure and therefore reduce the risk of sample photoinhibition during measurements. For the same reason, the blue laser (488 nm) intensity was set at the minimum value (0.3% - 220 μJ.cm^-2^) at which PSII fluorescence reached Fm to avoid excess excitation. Chl fluorescence was collected between 650-700 nm.

To estimate maximum photosynthetic capacity (Y), a series of 10 consecutive images was acquired (experimental time 6.33 s). A saturating pulse was provided by the external LED source after the fourth acquisition (Figure S1A). In the case of NPQ measurement, appropriate Z-stacks (xyzt experiment) were selected to include entire cells and maintain plastids in the acquisition field. Using the “sum slices” routine of Fiji, we transform xyzt files into xyt to calculate a time course of fluorescence changes. The delay for two consecutive acquisitions was set to 1 min to prevent too much laser exposure. 3 points were acquired for dark adapted samples, 6 during exposure to actinic red light and 6 points again in the dark to follow NPQ relaxation (experimental time 15 min, Figure S1B).

To calculate the ΦPSII parameter, lower intensities of the actinic laser were chosen, in order not to reach Fm, but rather a steady state level Fs. Maximum fluorescence Fm was instead achieved when the laser and the saturating Orange LED were switched on simultaneously.

In Fig. 1C and Figure S2, Chl fluorescence was instead measured with a conventional imaging setup (Speedzen, JBeamBio, France) described e.g. in (Seydoux et al., 2022).

### PSII yield and NPQ calculation from confocal images

Experimental files were imported to Fiji (Schindelin et al., 2012) and different Z acquisition were stacked to single images using Z project method and ‘sum slices’ function. Regions Of Interest (ROIs) were drawn on the Z project image to select single chloroplasts, cells or whole tissues or non-fluorescent regions that were used to estimate the background level. ‘Mean gray value’ of fluorescence was quantified for each single ROI and time. Background was subtracted to raw values for each time point. In the case of PSII capacity (Y): F’ value is the mean value for dark adapted samples measured with non-saturating laser light. Fm is instead the maximum value of the fluorescence obtained during exposition to a saturating pulse (Figure S1A). For NPQ, Fm is the mean value for dark adapted plant exposed to saturating laser intensity; Fm’ is the maximum fluorescence value samples exposed to actinic light or during the dark relaxation time (Figure S1B). In the case of radiolarians (Fig 3), Fs is the fluorescence measured in the presence of the non-saturating blue laser, while Fm is the fluorescence value achieved in the presence of the laser plus the saturating value of the orange LED.

### 3D reconstruction and fluorescence integration

Image processing was done adapting a pipeline previously developed for 3D reconstruction based on electron microscopy stacks (Uwizeye, 2021). Briefly, confocal images were pre-processed using a non-linear median filter (from Fiji) that preserves the edges. This is an essential prerequisite to calculate object volumes (see e.g. Fig 3B). We used 3DSlicer to perform semi-automatic segmentation. To calculate the volume of a reconstructed 3D model, we multiplied the number of voxels (volumetric picture elements) in the object by the size of the voxel : VI = (number of voxels in the object I) × (the size of the voxel Vi)

To calculate the fluorescence of each object in a confocal image, images were segmented to obtain the location of a given ROI). Data were used to calculate the sum of the voxel values of a given ROI, and therefore the mean fluorescence value of a given plastid or cell. In the case of *Phaeocystis cordata* photosymbiontic cells, where single plastid could not be imaged because of their small size, fluorescence parameters were calculated on cell sections, which we approximated with a circle (red circle in Fig 3E, F). To assess possible heterogeneous responses, we scanned fluorescence values along the radius. We then interpolated the corresponding fluorescence intensity to calculate fluorescence parameters (Fm, Fm’), and therefore NPQ, as a function of the angle. At least ten sections (i.e. 10 μm) were scanned for every cells to assess reproducibility.

### Principal components analysis (PCA)

The principal components analysis (PCA) is an exploratory technique that is used both to describe the structure of high dimensional data by reducing its dimensionality and to detect any groupings in the data set. It is a linear transformation that converts n original variables into n new variables (the principal components), which (i) are ordered by the amount of data variance explained by the component (ii) are uncorrelated and (iii) explain all variation in the data when sum up on the 6^th^ components. We performed PCA considering six observed variables: NPQav, NPQmax, Decay, Induction, Fmax and Area Fraction. The variables used here were calculated in automatized way from NPQ data. NPQav is the mean NPQ when a cell is exposed to light while NPQmax is the maximum of NPQ reached during the light exposition. Decay and Induction are the relaxation and induction rate of NPQ evaluated from the slope of NPQ changes during the first 2 minutes of the light to dark and the dark to light transitions, respectively. Area Fraction is the area percentage occupied by the chloroplast inside a cell. Fmax is the “Mean gray value” of fluorescence of a given cell chloroplasts for dark adapted samples (Fm).

Variables were measured in 175 cells from *P. patens* chloronema and caulonema in the WT or the *lhcsr1-2* KO strains. The type of cells (mutant/WT or chloronema/caulonema) did not play a role in the determination of the component and that they are used after to characterise the possible biological role of the components. All data are normalized by subtraction of the mean and division by the standard deviation so that the singular values decomposition is done on the correlation matrix of the data (Supplementary Table 1).

To represent the distribution of these normalized dimensional data for the 175 images, the direction (a 6-dimensional vector) giving the largest possible variance of the distribution was selected as the direction for the first principal component. Then, we selected the direction orthogonal to the previous one(s) giving the largest possible variance of the distribution as the direction for the second principal component. By repeating this procedure automatically, we identify vectors representing the scatter of the distribution from major ones to minor ones (Supplementary Table 2). Based on singular values decomposition, PCA is a principal axis rotation of the original variables that preserves the variation in the data. Therefore, the total variance of the original variables is equal to the total variance of the principal components. The principal component coefficients correspond to the percentage of explained variance. Statistical analysis was done with the R software (http://www.R-project.org). The table of the original observed variables used to construct the six components (Supplementary Table 3) provides the interpretation of the components.

## Supporting information

Supplemental Informations

## Acknowledgments

The authors thank prof. Alessandro Alboresi (University of Padova) for providing *P. patens* samples, dr. Charlotte Lekieffre (CEA Grenoble) for technical help with radiolarians, and the institutes that supported the collection of samples: EMBRC-France and the Laboratoire d’Océanographie de Villefranche-sur-Mer.

This project received funding from the European Research Council: ERC Chloro-mito (grant no. 833184) to F.C., J.D. M.S. and G.F. and the European Union H2020 Project BIOTEC-02-2019 GAIN4CROPS (grant no 862087) to G.C., D.T. and G.F.. Research was also supported by a Défi X-Life grant from CNRS to G.F. The ANR-11-BTBR-0008 Océanomics; ANR-15-IDEX-02 GlycoAlps and “Origin Of Life” Cross Disciplinary Projects of the Univ. Grenoble-Alpes; and ANR-17-EURE-0003 to O.B.. J.D. and F.C. were supported by CNRS and ATIP-Avenir program. Funding by the LabEx GRAL (ANR-10-LABX-49-01), financed within the University Grenoble Alpes graduate school (Ecoles Universitaires de Recherche) CBH-EUR-GS (ANR-17-EURE-0003) is also acknowledged.

## Author contributions

Conceptualization, G.F.; Methodology, M.S., D.B., D.T. and G.F.; Image analysis C.U., and H.H.; Data Analysis, M.S. and O.B.; Investigation, M.S., H.H., D.T. and G.F. Resources, D.Y., F.C., C.G. and D.T.; Writing-Original Draft, M.S., D.T. and G.F.; Writing-Review & Editing, M.S., G.C., D.T. and G.F.; Funding Acquisition, G.F.; Supervision, G.C., J.D., D.T. and G.F.

## Figures

**Supplementary Video 1 (related to Figure 2): Plastids movements inside a *P. patens* filament**. Overlapped bright field (grey) and chlorophyll fluorescence (red) images were acquired with an upright Leica DM6 B (HC PL FLUOTAR 20x/0.50 objective, Excitation filter 450 nm) optical microscope. Image size= 342.58 × 256.89 μm. Time series were acquired with a delay of 30 s between different images and reproduced at 2 frames per second.

**Supplementary Videos 2-5 (related to Figure 1): Time course of fluorescence changes in *P. patens***. Time course Z-projections (15-20 slices) of *P. patens* protonema exposed for 6 min to 500 μmols photons m^-2^ s^-1^. Time series that acquired with 1 min delay are reproduced at 1 frame per second (fps). The sum of raw fluorescence values for the different stacks is represented in a false-colour scale bar on the top.

