## Supplemental Informations for "Tailoring confocal microscopy for in-cell photophysiology studies"

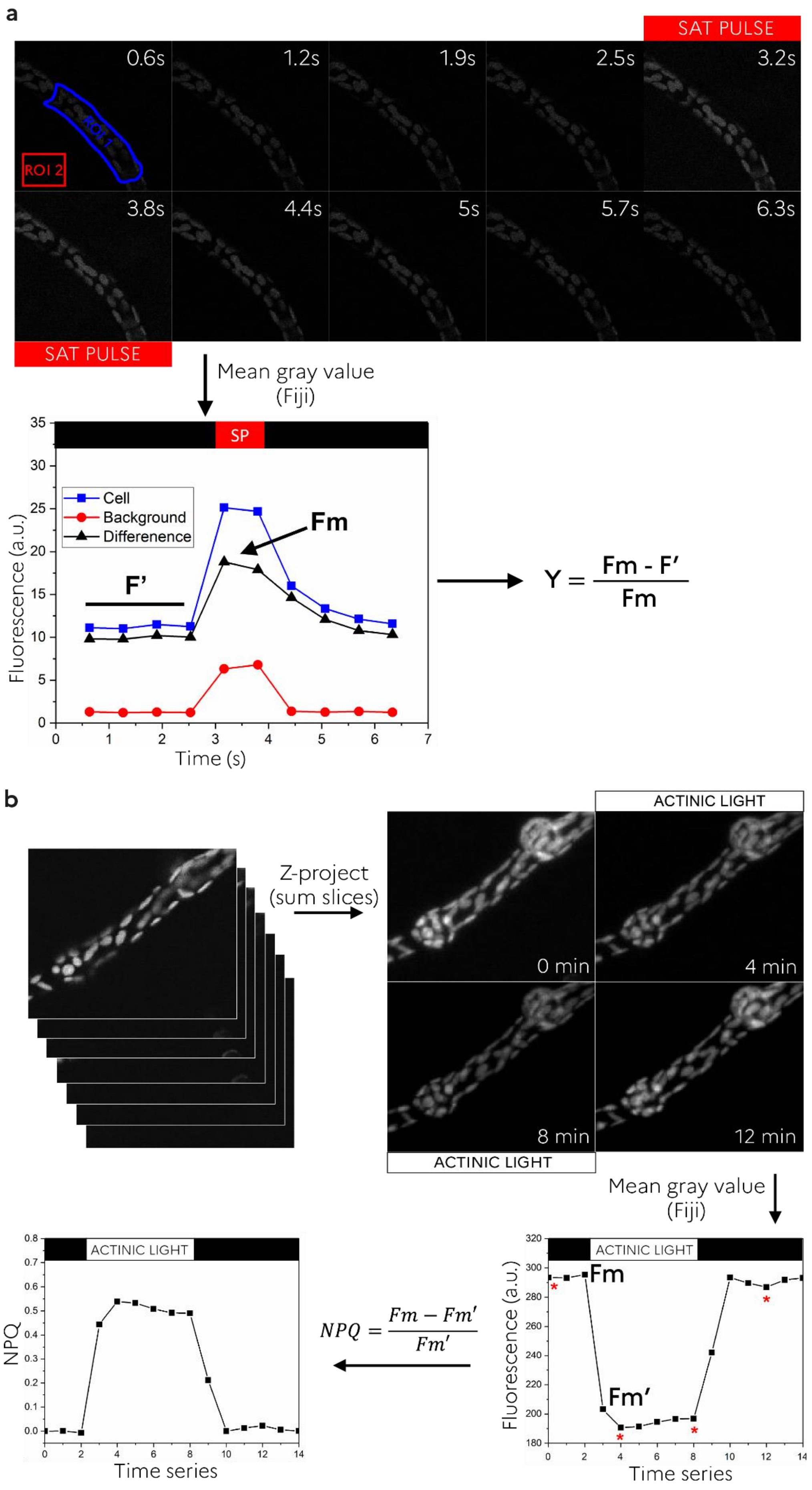
a
SAT PULSE
0.6s
1.2s
1.9s
2.5s
3.2s
ROI 2
3.8s
4.4s
5s
5.7s
6.3s
SAT PULSE
Mean gray value (Fiji)
SP
Cell
Background
Differenence
Fm
F'
Fluorescence (a.u.)
Time (s)
Y = Fm - F' / Fm
b
Z-project (sum slices)
ACTINIC LIGHT
0 min
4 min
8 min
12 min
ACTINIC LIGHT
Mean gray value (Fiji)
ACTINIC LIGHT
NPQ
Time series
NPQ = Fm − Fm' / Fm'
ACTINIC LIGHT
Fm
Fm'
Fluorescence (a.u.)
Time series

**Figure S1 (related to Figure 1): Analysis of confocal images with Fiji.** (A) To measure the 'Y' parameters (PSII yield) we calculated the "mean gray value" of ROIs (ROI 1, blue: *P. patens* cell; ROI 2, red background). (B), Background fluorescence (red), which is altered by external light source (Saturating pulse, SP, red bar) was subtracted from the sample fluorescence (blue) value to obtain the real fluorescence values (difference, black). By plotting the results of this calculation for sequential acquisitions, we obtained a time series, which we used to calculate Y values. For NPQ experiments, xyzt files (C) were converted to xyt files (D) using the "sum slices" function of Fiji. Images were analysed as in (A) to calculate NPQ (E). Fm is the fluorescence value of the dark-adapted sample (top black bar) while Fm' is the fluorescence value measured during exposition to actinic light (top white bar). Both values are needed to evaluate NPQ. Red asterisks in (F) represent the fluorescence values calculated from the images represented in panel B, right.

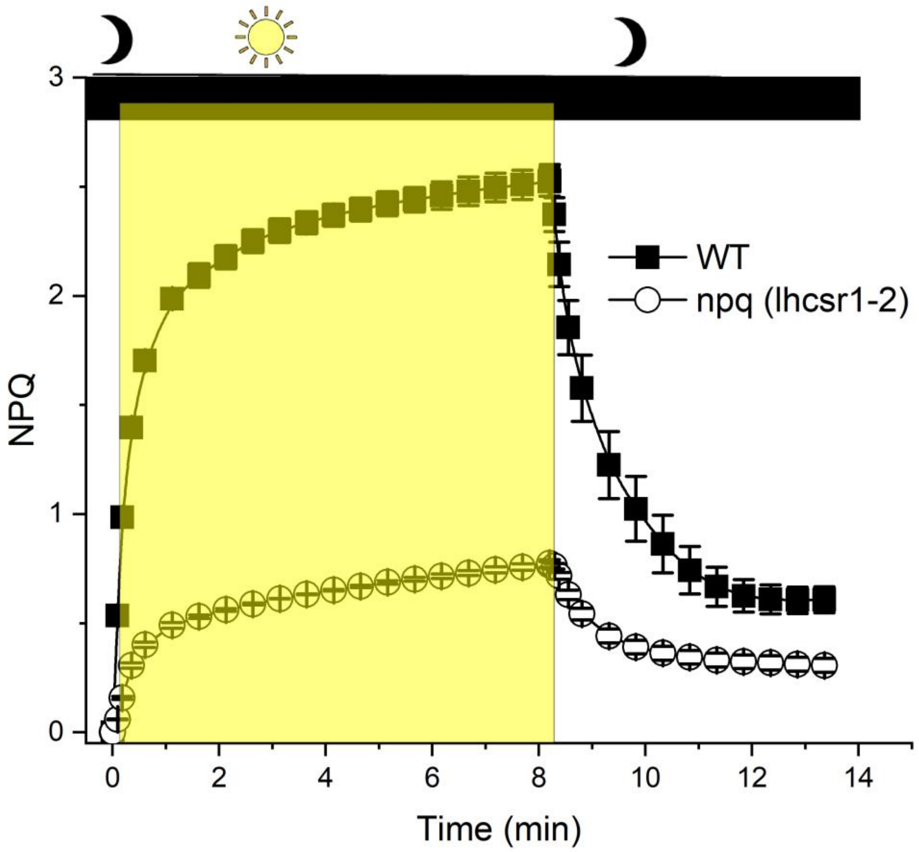


**Figure S2 (related to Figure 1): NPQ in *P. patens*, measured with a chlorophyll fluorescence imaging camera.** WT (black squares) and *lhcsr1/2 (empty circles) P. patens* protonema were exposed to saturating light intensity (yellow overlay, 500 µmols photons $m^{-2}$ $s^{-1}$) for 8 min before measuring photosynthetic parameters using a speedzen Chl fluorescence imaging setup (JBeambio, France) (Seydoux et al., 2022). Same symbols and box colours as in Fig 1F.

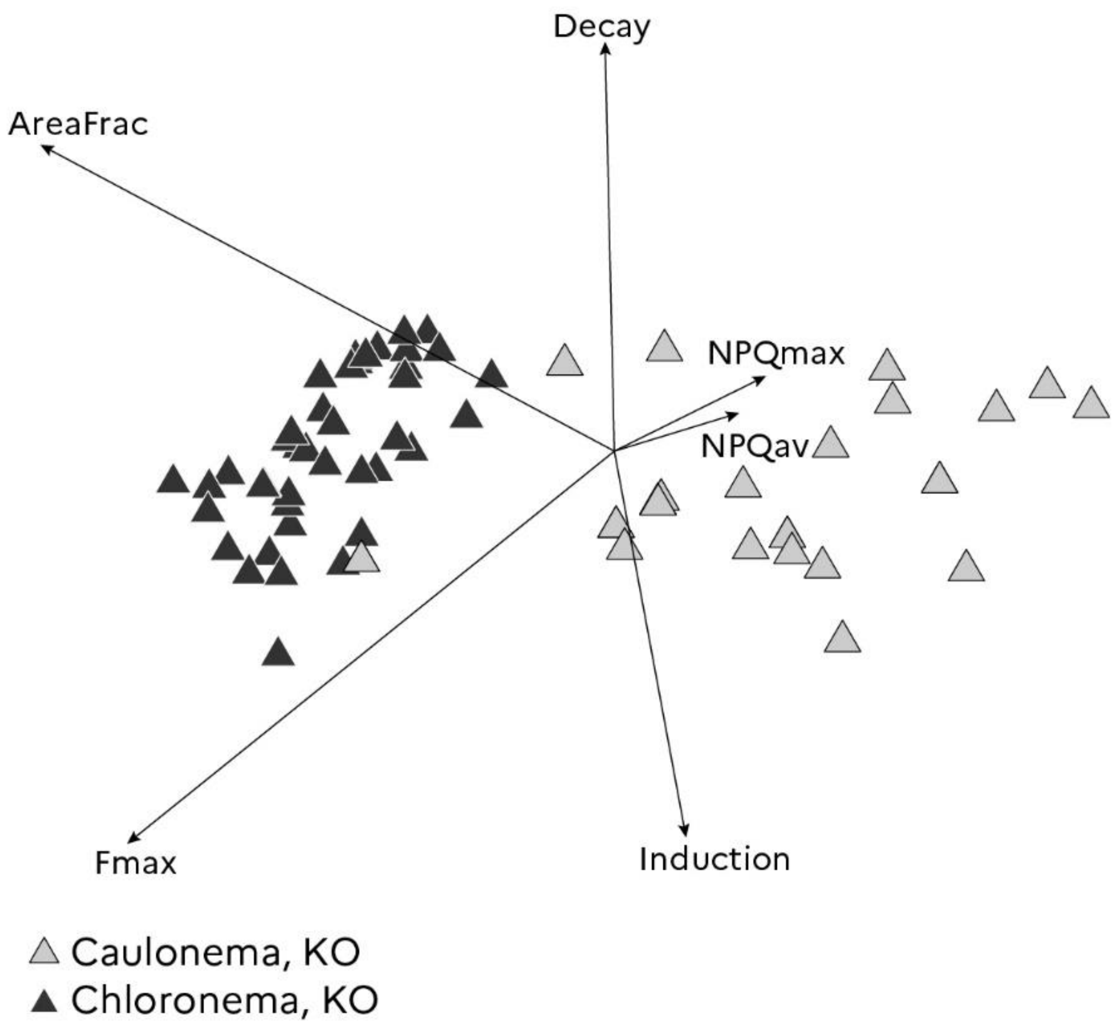


**Figure S3 (related to Figure 2): PCA analysis of NPQ features in the *P. patens lhcsr1/2* KO-mutant: second and third components.** Second and third components of a PCA realised on 63 P. patens lhcrs1/2 KO cells (solid symbols chloronema, open symbols: caulonema). The first two components represent roughly 88% of the variance and the first three components represents more than 94% of the variance (Supplementary Table 2).

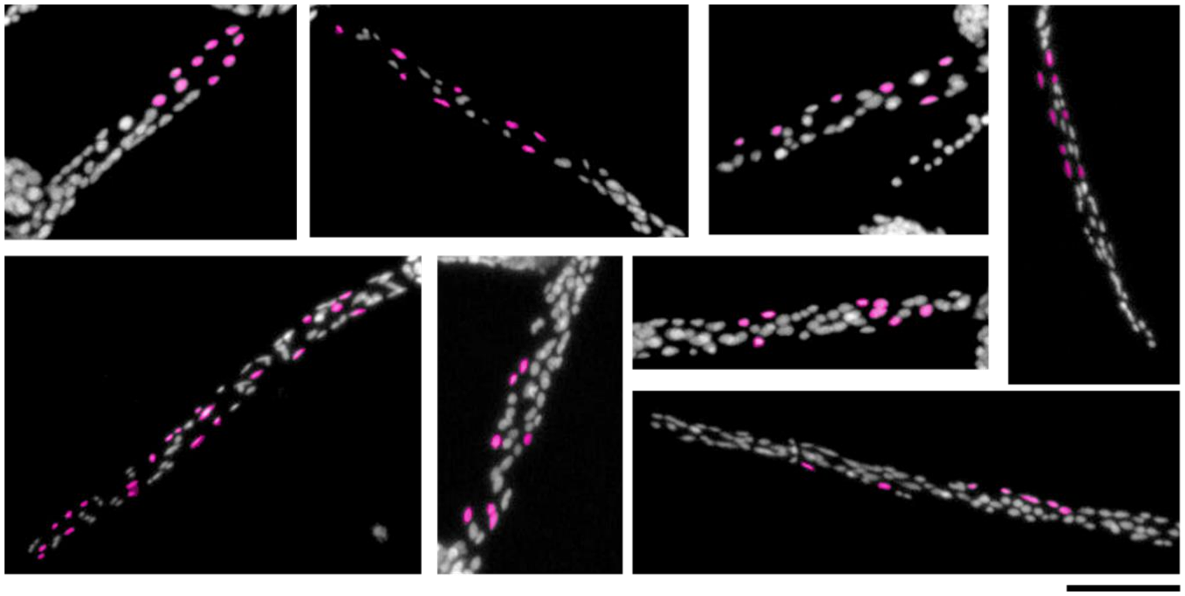

**Figure S4 (related to Figure 2): NPQ analysis of *P. Patens* single chloroplasts.** Among the numerous plastids of *P. patens* protonema, some single chloroplasts (highlighted in pink) were selected because they never overlap with each other during the confocal microscope acquisition. Chloroplasts movements were tracked manually and fluorescence values were calculated in the dark and after 6 min of light exposure (500 µmols photons $m^{-2}$ $s^{-1}$). Scale bar: 50 µm.

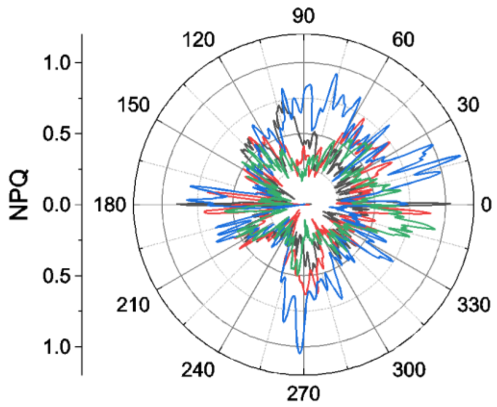

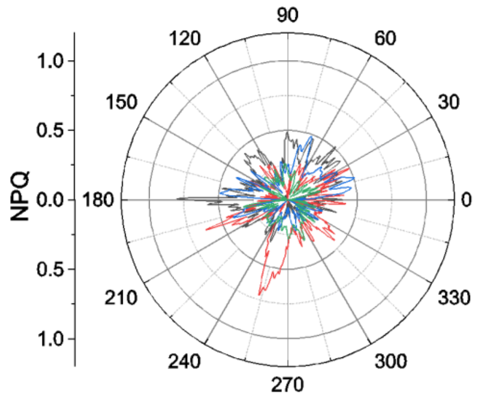


**Figure S5 (related to Figure 3): subcellular analysis of NPQ in radiolarians.** radar plot of the NPQ in large (A) and small (B) symbiotic microalgae. NPQ was calculated from images acquired in the dark and after exposure to actinic light (500 µmols photons $m^{-2}$ $s^{-1}$) for 15 minutes. Different colours represent different radiolarian cells

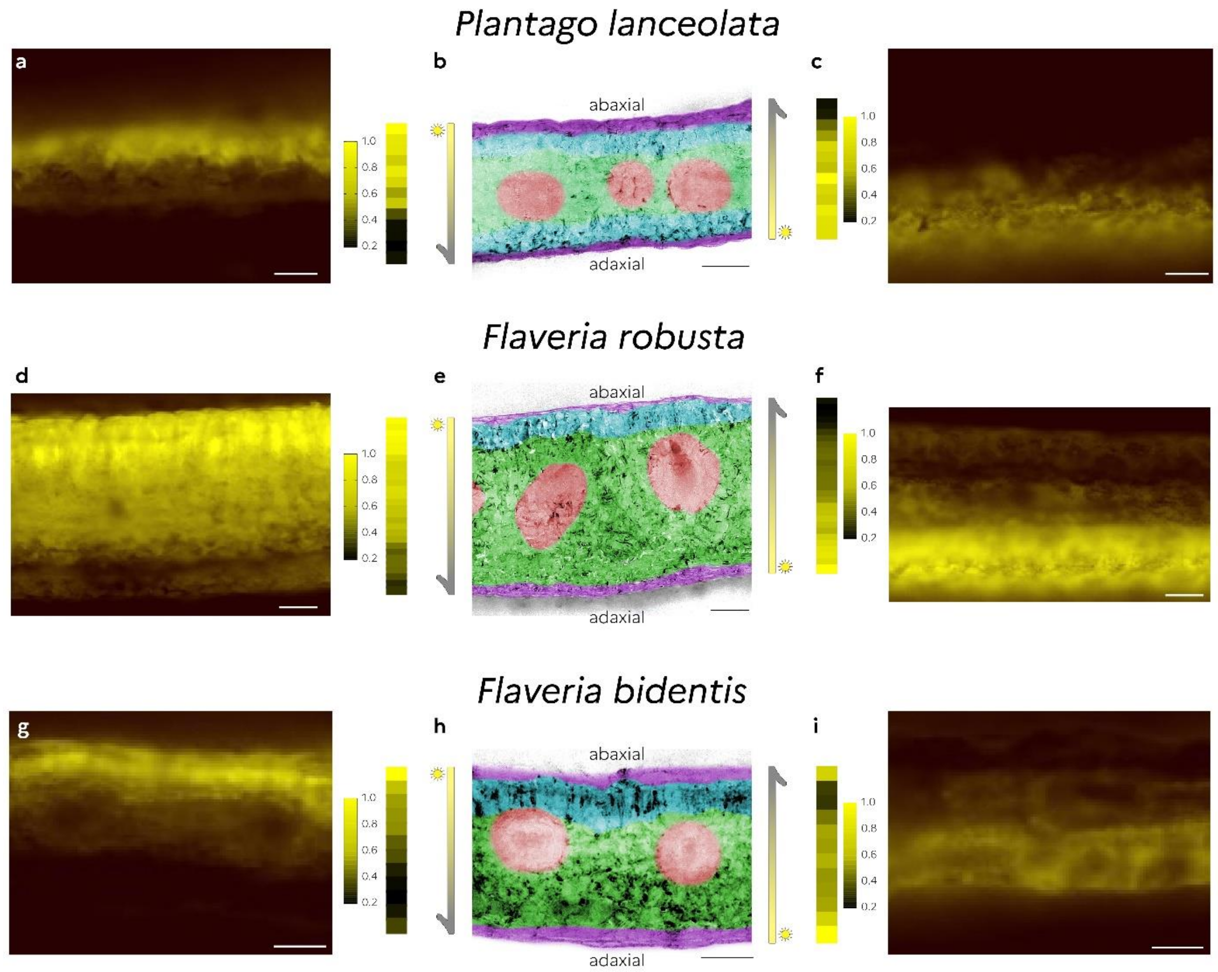


**Figure S6 (related to Figure 4): Light gradients inside leaves**. Light gradients are visualised by the difference in chlorophyll fluorescence (figured as yellow in false colour) intensity upon illumination in the adaxial to abaxial side (bottom to top –panels A, D and G) and adaxial to abaxial (bottom to top – panels C, F and I) for a representative cross section of leaves with different anatomies (panels B, E and H). The following colours were used to highlight the different leaf tissues: purple: epidermis; blue: palisade parenchyma; green: spongy parenchyma; red: vascular tissue

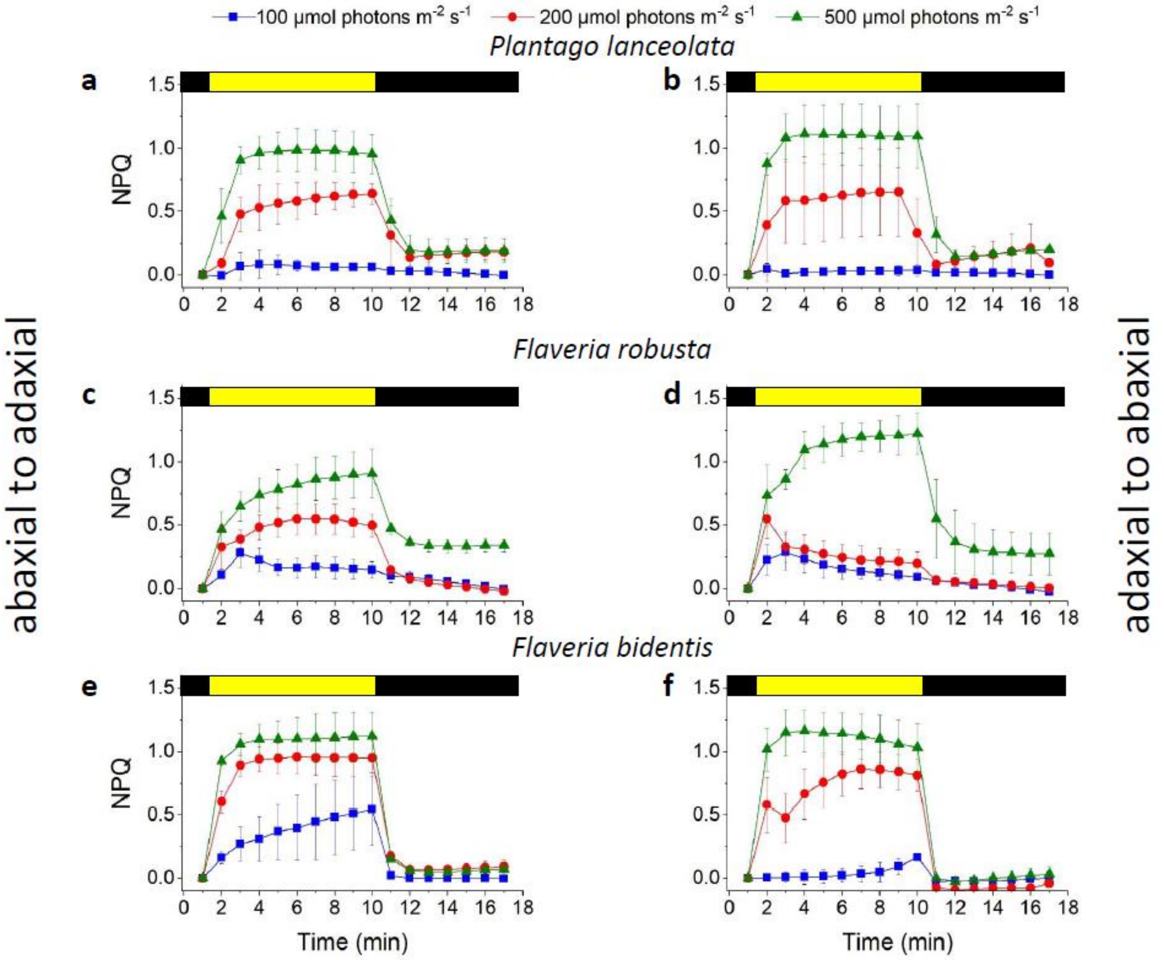


**Figure S7 (related to Figure 4): NPQ features are modulated by leaves architectures**. NPQ measured upon illumination in the adaxial to abaxial side (top to bottom –panels A, C and E) and abaxial to adaxial (bottom to top – panels B, D and F) for three leaves with different anatomies as presented in Fig. 4. Shown are means ± SDs (n=3)

| | Comp. 1 | Comp. 2 | Comp. 3 | Comp. 4 | Comp. 5 | Comp. 6 |
|---|---|---|---|---|---|---|
| Standard deviation | 1.9182243 | 1.2630922 | 0.60785746 | 0.45836413 | 0.35531842 | 0.138470303 |
| Proportion of Variance | 0.6132641 | 0.2659003 | 0.06158178 | 0.03501628 | 0.02104186 | 0.003195671 |
| Cumulative Proportion | 0.6132641 | 0.8791644 | 0.94074619 | 0.97576247 | 0.99680433 | 1.000000000 |

**Supplementary Table 1 (related to Figure 2): Correlation table of the six observed variables**

| | NPQmax | NPQav | Decay | Induction | AreaFrac | Fmax |
|---|---|---|---|---|---|---|
| NPQmax | 1.00000000 | 0.97681818 | 0.8537735 | 0.812678625 | - 0.070494246 | 0.2514870 |
| NPQav | 0.97681818 | 1.00000000 | 0.8392191 | 0.837617127 | - 0.020657062 | 0.2833078 |
| Decay | 0.85377353 | 0.83921910 | 1.0000000 | 0.701930935 | 0.195843670 | 0.3648589 |
| Induction | 0.81267862 | 0.83761713 | 0.7019309 | 1.000000000 | 0.009117372 | 0.3559048 |
| AreaFrac | -0.07049425 | -0.02065706 | 0.1958437 | 0.009117372 | 1.000000000 | 0.6719592 |
| Fmax | 0.25148703 | 0.28330778 | 0.3648589 | 0.355904772 | 0.671959243 | 1.0000000 |

**Supplementary Table 2 (related to Figure 2): Principal Component Analysis results.**

| | 1st Comp | 2nd Comp | 3rd Comp | 4th Comp | 5th Comp | 6th Comp |
|---|---|---|---|---|---|---|
| NPQmax | 24.4496826 | 3.7675641 | 0.9885236 | 3.6558842 | 14.9788645 | 52.15948102 |
| NPQav | 24.8978544 | 2.5525258 | 0.2549017 | 0.2827022 | 25.9588445 | 46.05317140 |
| Decay | 22.5153461 | 0.0140642 | 29.3140005 | 3.7408834 | 43.5351179 | 0.88058787 |
| Induction | 21.4494317 | 0.8431956 | 26.1007806 | 42.9901977 | 8.2970322 | 0.31936221 |
| AreaFrac | 0.6559251 | 53.9600310 | 16.4085692 | 21.8731751 | 6.5749696 | 0.52732993 |
| Fmax | 6.0317601 | 38.8626193 | 26.9332244 | 27.4571575 | 0.6551713 | 0.06006757 |

**Supplementary Table 3 (related to Figure 2): Contribution table of the six observed variables to the construction of the six components.** For example, the first component is constructed with a contribution of NPQmax with 24.5 %, NPQav with 24.9%, Decay with 22.5% and Induction with 21.5%

Seydoux, C., Storti, M., Giovagnetti, V., Matuszyńska, A., Guglielmino, E., Zhao, X., Giustini, C., Pan, Y., Blommaert, L., Angulo, J., Ruban, A. v., Hu, H., Bailleul, B., Courtois, F., Allorent, G., & Finazzi, G. (2022). Impaired photoprotection in *Phaeodactylum tricornutum* KEA3 mutants reveals the proton regulatory circuit of diatoms light acclimation. *New Phytologist*, *234*(2), 578–591. https://doi.org/10.1111/nph.18003
